## Supplementary material for "Lipid Peroxidation and Type I Interferon Coupling Fuels Pathogenic Macrophage Activation Causing Tuberculosis Susceptibility": Key Resources Table

| **REAGENT or RESOURCE** | **SOURCE** | **IDENTIFIER** |
| --- | --- | --- |
| **Antibodies** | | |
| Anti-iNOS antibody | Abcam | Cat# ab15323, RRID:AB_301857 |
| Iba1/AIF-1 (E4O4W) XP® Rabbit mAb | Cell Signaling Technology | Cat# 17198, RRID:AB_2820254 |
| GFP Polyclonal Antibody | Invitrogen | Cat#A11122, RRID:AB_221569 |
| IFNAR1 Monoclonal Antibody (MAR1-5A3), Functional Grade | Thermo Fisher Scientific | Cat# 16594585,  RRID:AB_1210688 |
| Mouse IgG1 kappa Isotype Control Antibody (P3.6.2.8.1) | Thermo Fisher Scientific | Cat#14471482,  RRID:AB_470111 |
| TNF alpha Monoclonal Antibody (XT22) | Thermo Fisher Scientific | Cat#MM350D,  RRID:AB_223528 |
| KEAP1 Rabbit anti-Human, Mouse, Rat, Polyclonal Antibody | Proteintech Group Inc | Cat#10503-2-AP  RRID:AB_2132625 |
| NRF2/NFE2L2 Rabbit anti-Human, Mouse, Rat, Polyclonal Antibody | Proteintech Group Inc | Cat#16396-1-AP  RRID:AB_2782956 |
| Anti-βTrCP Ab | Proteintech Group Inc | Cat#28393-1-AP  RRID:AB_2935467 |
| BACH1 Polyclonal Antibody | Invitrogen | Cat#PA5-117013  RRID:AB_2901643 |
| Anti-Histone H3 Ab | Cell Signaling Technology | Cat#9715s  RRID:AB_331563 |
| Anti-β-tubulin Ab | Santa Cruz Biotechnology | Cat#sc-55529  RRID:AB_2210962 |
| Anti-p-cJun Ab | Cell Signaling Technology | Cat#9261s  RRID:AB_2130162 |
| Anti-Myc Ab | Santa Cruz Biotechnology | Cat# sc-788  RRID:AB_631277 |
| Recombinant Anti-c-Myc antibody [Y69] | Abcam | Cat# ab32072  RRID:AB_731658 |
| Recombinant Anti-Ferritin Heavy Chain antibody [EPR18878] | Abcam | Cat# ab183781,  RRID:AB_2940987 |
| Recombinant Anti-Ferritin Light Chain antibody [EPR5260] | Abcam | Cat#ab109373  RRID:AB_1086271 |
| Anti-4 Hydroxynonenal antibody | Abcam | Cat# ab46545,  RRID:AB_722490 |
| Anti-NRF1 Ab | Santa Cruz Biotechnology | Cat#sc-515360, |
| Anti-GPX1 Ab | Proteintech Group Inc | Cat#29329-1-AP  RRID:AB_2918283 |
| Anti-GPX4 Ab | Proteintech Group Inc | Cat#67763-1-Ig  RRID:AB_2909469 |
| Anti-mouse IgG, HRP-linked Ab | Cell Signaling Technology | Cat#7076s  RRID:AB_330924 |
| Anti-rabbit IgG, HRP-linked Ab | Cell Signaling Technology | Cat#7074s  RRID:AB_2099233 |
| ImmPRESS HRP Anti-Rabbit IgG (Peroxidase) Polymer Detection Kit, made in Goat | Vector Laboratories | Cat#MP-7451, RRID:AB_2631198 |
| ImmPRESS™ HRP Anti-Mouse IgG (Peroxidase) Polymer Detection Kit, made in Goat | Vector Laboratories | Cat#MP-7452, RRID:AB_2744550 |
| Goat anti-Rabbit IgG (H+L) Cross-Adsorbed Secondary Antibody, Alexa Fluor™ 546 | Invitrogen | Cat# A-11010,  RRID:AB_2534077 |
| Goat anti-Rabbit IgG (H+L) Highly Cross-Adsorbed Secondary Antibody, Alexa Fluor™ Plus 647 | Invitrogen | Cat# A-32733,  RRID:AB_2633282 |
| Iba1 Antibody (1022-5), Alexa Fluor® 647 | Santa Cruz Biotechnology | Cat# sc-32725,  RRID:AB_667733 |
| ChromoMap DAB Kit | Roche | Cat#760-159 |
| HRP/DAB detection kit | Abcam | Cat# ab64261 |
| Tris based buffer-Cell Conditioning 1 (CC1) | Roche | Cat#950-124 |
| **Bacterial and virus strains** | | |
| *Mycobacterium tuberculosis* H37Rv | ATCC | Cat# 27294 |
| *Mycobacterium bovis* BCG | ATCC | Cat# 35737 |
| Erdman(SSB-GFP, *smyc′*::mCherry) | Lavin et al., 2022 | N/A |
| **Biological samples** |  |  |
| **Chemicals, peptides, and recombinant proteins** | | |
| Pexidartinib (PLX3397) | Selleckchem | Cat# S7818 |
| Sotuletinib (BLZ945) | MedChemExpress | Cat# HY-12768 |
| GW-2580 | MedCHemExpress | Cat#HY-10917 |
| 10058-F4 | Selleckchem | Cat# S7153 |
| Ferrostatin-1 | Selleckchem | Cat#S7243 |
| D-JNK-1 | MedChemExpress | Cat# HY-P0069 |
| Deferoxamine mesylate (DFOM) | Sigma Aldrich | Cat#D9533 |
| Butylated hydroxyanisole (BHA) | Sigma Aldrich | Cat# B1253 |
| Hygromycin B | Roche | Cat# 10843555001 |
| DMEM/Ham's F-12 50/50 Mix [+] L-glutamine | Corning® | Cat# 45000-344 |
| DMEM, 1X (Dulbecco’s Modified Eagle’s Medium) | Corning® | Cat# 10-013-CV |
| Fetal Bovine Serum (FBS) | Hyclone^TM^, | Cat# SH30071.03 |
| Hoechst 33342 | Fisher Scientific | Cat# H3570 |
| Paraformaldehyde Solution 4% in PBS | Fisher Scientific | Cat# J19943-K2 |
| L-Glutamine | Corning® | Cat# 25-005-CI |
| Penicillin Streptomycin solution | Corning® | Cat# 30-002-CI |
| HEPES buffer | Corning® | Cat# 25-060-CI |
| L929 Cell Conditioned Media (LCCM) | This paper | N/A |
| Murine IFN-gamma | Peprotech | Cat# 315-05 |
| Murine Interleukin -3 | Peprotech | Cat# 213-13 |
| Murine Interleukin-4 | Peprotech | Cat# 214-14 |
| Murine TNF-alpha | Peprotech | Cat# 315-01A |
| Lymphoprep^TM^ (1.077A) | STEMCELL | Cat#07801 |
| Poly Ethylene Glycol (PEG), Bioultra-8000 | Sigma | Cat#89510 |
| 5M NaCl | Invitrogen | Cat#AM9759 |
| Tris Hydrochloride, 1M solutions (pH 8.0) | Fisher Scientific | Cat#77-86-1 |
| Ultrapure 0.5 M EDTA pH 8.0 | Invitrogen | Cat#15575-038 |
| Ambion^TM^ Nuclease-free Water | Invitrogen | Cat#AM9932 |
| SpeedBead Magnetic Carboxylate Modified Particles | GE Healthcare | Cat#65152105050250 |
| DynaMag^TM^-96 side | Life Technologies^TM^ | Cat#12331D |
| Glycine | Sigma | Cat#50046 |
| NaOH Solution | Sigma | Cat#72068 |
| Proteinase K | Ambion | Cat#AM2546 |
| Middlebrook 7H9 Broth | BD Biosciences | Cat# 271310 |
| Middlebrook 7H10 Agar | BD Biosciences | Cat# 262710 |
| Rapiclear 1.47 | SunJin Lab Co. | Cat# NC1660944 |
| ProLong™ Gold Antifade Mountant | Invitrogen™ | Cat# P36934 |
| Cycloheximide | Cell Signaling Technology | Cat#2112 |
| **Critical commercial assays** | | |
| Live-or-Dye™ 594/614 Fixable Viability Staining Kits | Biotium | Cat# 32006 |
| TaqMan™ Environmental Master Mix 2.0 | Fisher Scientific | Cat#4396838-5mL |
| Lipid Peroxidation (MDA) Assay Kit (Colorimetric/Fluorometric) | Abcam | Cat# ab118970 |
| BODIPY™ 581/591 C11 (Lipid Peroxidation Sensor) | Thermo Fisher Scientific | Cat# D3861 |
| CellROX™ Green Reagent | Thermo Fisher Scientific | Cat# C10444 |
| Click-iT™ Lipid Peroxidation Imaging Kit | Thermo Fisher Scientific | Cat# C10446 |
| Nuclear Extraction Kit | Signosis | Cat# SK-0001 |
| NRF2(ARE) EMSA Kit | Signosis | Cat# GS-0031 |
| HCR^TM^ IFNβ probe set | Molecular instruments | N/A |
| HCR^TM^ Buffers | Molecular instruments | N/A |
| Antioxidant Assay Kit | Cayman chemical | Cat# 709001 |
| RNeasy plus mini kit | Qiagen | Cat#74136 |
| Invitrogen™ SuperScript™ III First-Strand Synthesis SuperMix | Invitrogen | Cat#18080400 |
| GoTaq qPCR Mastermix | Promega | Cat#A6002 |
| PAXgene Blood RNA kit | Qiagen, Hilden, Germany | Cat #762164 |
| SureSelect Strand-Specific mRNA Library Prep kit | Agilent, Santa Clara, USA | Cat #5190–6411 |
| **Deposited data** | | |
| **Experimental models: Cell lines** | | |
| **Experimental models: Organisms/strains** | | |
| Mouse: C57BL/6J | The Jackson Laboratory | Stock No.: 000664, RRID:IMSR_JAX:000664 |
| Mouse: B6J.C3-*Sst1^C3HeB/Fej^*Krmn | Pichugin et al., 2009 | Stock No: 043908-UNC  <https://www.mmrrc.org> |
| Mouse: C3HeB/FeJ | The Jackson Laboratory | Stock No.: 000658, RRID:IMSR_JAX:000658 |
| Mouse: (C3HeB/FeJ X B6.Sst1S)F1 | This study | N/A |
| Mouse: B6.Sst1S,*ifnb1*-YFP | This study/ Rosenbloom et al, 2022, Yabaji et al., 2023 | N/A |
| **Oligonucleotides** | | |
| Mtb specific FP: GGAAATGTCACGTCCATTCATTC | Yabaji et al., 2022 | N/A |
| Mtb specific RP: GCGTTGTTCAGCTCGGTA |  |  |
| Mtb specific probe: 56-FAM/AGCTTGGTCAGGGACTGCTTCC/36-TAMSp/ |  |  |
| BCG specific FP: GTGGTGGAGCGGATTTGA |  |  |
| BCG specific RP: CAACCGGACGGTGATCC |  |  |
| BCG specific probe: /5Cy5/TTCTGGTCG/TAO/ACGATTGGCACATCC/3IAbRQSp/ |  |  |
| Primers  *Ifnb, Rsad2, Trib3, Chac1, b-actin, 18S* | Bhattacharya et al., 2021 | N/A |
| *Fth*  FP: 5’-TGTATGCCTCCTACGTCTATCT-3'  RP: 5’-CCTCATGAGATTGGTGGAGAAA-3' | This study | N/A |
| *Ftl*  FP: 5’-AGGAGGTGAAACTCATCAAGAA-3' RP: 5’-TGAGGCGCTCAAAGAGATAC-3' | This study | N/A |
| *Myc*  FP: 5’-TCTCCACTCACCAGCACAACTACG-3'  RP: 5’-ATCTGCTTCAGGACCCT-3' | This study | N/A |
| *Hmox-1*  FP: 5’- CCTTCCCGAACATCGACAGCC- 3'  RP: 5'- GCAGCTCCTCAAACAGCTCAA- 3' | This study | N/A |
| *Nqo1*  FP: 5'- CCTCGCTGGAAAAAGAAGTG- 3'  RP: 5'- GGAGAGGATGCTGCTGAAAG- 3' | This study | N/A |
| *Nfe2l2*  FP: 5'-CCTCGCTGGAAAAAGAAGTG- 3'  RP:5’-GGAGAGGATGCTGCGGAAAG-3’ | This study | N/A |
| *Gpx1*  FP:5’-CACCAGGAGAATGGCAAGAA-3'  RP:5’-CATTCCGCAGGAAGGTAAAGA-3' | This study | N/A |
| *CIITA*  FP:5’-CTTCAAGCAGCCTCAGTATC-3'  RP:5’-ATGTGTCCTCTGTCTCATTTAC-3' | This study | N/A |
| **Recombinant DNA** | | |
| **Software and algorithms** | | |
| Graphpad Prism 9.5.1 (528) | Graphpad | <https://www.graphpad.com/>, RRID: SCR_002798 |
| Microsoft office | Microsoft | https://www.office.com/?auth=2 |
| Halo HighPlex FL v4.2.3 | Indica Labs Inc. | https://indicalab.com/halo/ |
| EndnoteX9 | Clarivate Analytics | https://endnote.com/downloads |
| Imaris Viewer | Oxford Instruments | <https://imaris.oxinst.com/microscopy-imaging-software-free-trial>?  source=viewer |
| ImageJ | Schneider et al., 2012 | <https://imagej.nih.gov/ij/>, SCR_003070 |
| STAR | STAR | RRID: SCR_004463 |
| featureCounts | featureCounts | RRID: SCR_012919 |
| DESeq2 | DESeq2 | RRID: SCR_015687 |
| limma | limma | RRID: SCR_010943 |
| GSEA | GSEA | RRID: SCR_003199 |
| Seurat | Seurat | RRID: SCR_007322 |
| Cytoscape | [Cytoscape](http://cytoscape.org/) | RRID: SCR_003032 |
| Trimmomatic | Trimmomatic | RRID: SCR_011848 |
| GEO | [GEO](https://www.ncbi.nlm.nih.gov/geo/) | RRID: SCR_005012 |
| GENIE3 | [GENIE3](http://www.genie3.org/) | RRID: SCR_000217 |
| RStudio | RStudio | RRID: SCR_000432 |
| Enrichr | Enrichr | RRID: SCR_001575 |
| **Other** | | |
| Operetta CLS HCA System | Operetta^TM^ | https://www.perkinelmer.com/in/lab-solutions/product/operetta-cls-system-hh16000020 |
| Vibratome | Leica VT1200 S | https://www.leicabiosystems.com/us/research/vibratomes/leica-vt1200/ |
| SP5 Confocal Microscope | Leica | N/A |
| LAS-4000 | FujiFilm | N/A |
| Automate in vivo manual gravity perfusion system for mice double 140 mL – IV 4140 | Braintree Scientific, Inc | Cat# IV 4140 |
